# The emergence of phase separation as an organizing principle in bacteria

**DOI:** 10.1101/2020.08.05.239012

**Authors:** C.A. Azaldegui, A.G. Vecchiarelli, J.S. Biteen

**Affiliations:** University of Michigan

## Abstract

Recent investigations in bacteria suggest that membraneless organelles play a crucial role in the subcellular organization of bacterial cells. However, the biochemical functions and assembly mechanisms of these compartments have not yet been completely characterized. This Review assesses the current methodologies used in the study of membraneless organelles in bacteria, highlights the limitations in determining the phase of complexes in cells that are typically an order of magnitude smaller than a eukaryotic cell, and identifies gaps in our current knowledge about the functional role of membraneless organelles in bacteria. Liquid-liquid phase separation (LLPS) is one proposed mechanism for membraneless organelle assembly. Overall, we outline the framework to evaluate LLPS *in vivo* in bacteria, we describe the bacterial systems with proposed LLPS activity, and we comment on the general role LLPS plays in bacteria and how it may regulate cellular function. Lastly, we provide an outlook for super-resolution microscopy and single-molecule tracking as tools to assess condensates in bacteria.

**Statement of Significance:** Though membraneless organelles appear to play a crucial role in the subcellular organization and regulation of bacterial cells, the biochemical functions and assembly mechanisms of these compartments have not yet been completely characterized. Furthermore, liquid-liquid phase separation (LLPS) is one proposed mechanism for membraneless organelle assembly, but it is difficult to determine subcellular phases in tiny bacterial cells. Thus, we outline the framework to evaluate LLPS *in vivo* in bacteria and we describe the bacterial systems with proposed LLPS activity in the context of these criteria.

## Introduction

Over the last decade, it has become clear that membraneless organelles play a key role in subcellular spatial organization, and a surge of studies indicates that these compartments are ubiquitous in eukaryotic cells (1). More surprisingly, recent investigations in bacteria suggest that membraneless organelles play a crucial role in the subcellular organization of bacterial cells as well (2–5). However, the biochemical functions and assembly mechanisms of these compartments have not yet been completely characterized. Our goal in this review is therefore to assess the current methodologies used in the study of membraneless organelles in bacteria, to highlight the limitations in determining the phase of complexes in cells that are typically an order of magnitude smaller than a eukaryotic cell, and finally, to identify gaps in our current knowledge about the functional role of membraneless organelles in bacteria.

Liquid-liquid phase separation (LLPS) is one proposed mechanism for membraneless organelle assembly. In this process, biomolecules separate in solution to form a condensed liquid phase with material properties distinct from those of the surrounding dilute phase. Observed as clusters, hubs, foci, puncta, or droplets, the membraneless organelles formed by this process are referred to as biomolecular condensates (condensates hereafter) (6–8). Condensates assemble through a collection of weak protein-nucleic acid and/or protein-protein interactions, typically involving the intrinsically disordered regions (IDRs) of their biomolecular building blocks (6, 9). For a more in-depth description of LLPS and how it drives condensate formation, we refer the reader to other reviews (3, 6, 10, 11). Certain condensate properties currently serve as criteria for determining if a membraneless compartment is indeed phase-separated. In the context of their studies of the nucleolus and P granules (12, 13), Hyman and colleagues proposed the following criteria for defining LLPS in eukaryotic cells (Fig 1A-C): nucleation of the condensate components into a spherical shape, the ability of the droplets to fuse, and sufficient mobility of the components to permit both rearrangement within the condensate and exchange across its boundary (10). Recently, two additional criteria have been proposed as LLPS indicators (Fig 1D-E): a concentration dependence to the molecular components and slowed diffusion across the condensate boundary (14, 15). Condensates form when their building blocks reach a saturation threshold, *c*_sat_, and the size of the droplet scales with the degree of supersaturation, leaving the surrounding cytoplasm buffered to *c*_sat_ (8, 16). Furthermore, their lack of a membrane allows condensates to tune their composition, size, and structure in real time, and enables them to dissolve, swap components, and transition between phase states. The properties described for eukaryotic cells provide an experimental framework to evaluate LLPS in live bacterial cells. However, a key hurdle specific to the study of membraneless organelles in bacteria is cell size. Many of the liquid-like features that are easily detected using traditional microscopy approaches on massive eukaryotic cells are beyond the resolution limit in bacteria.

**Figure 1.**
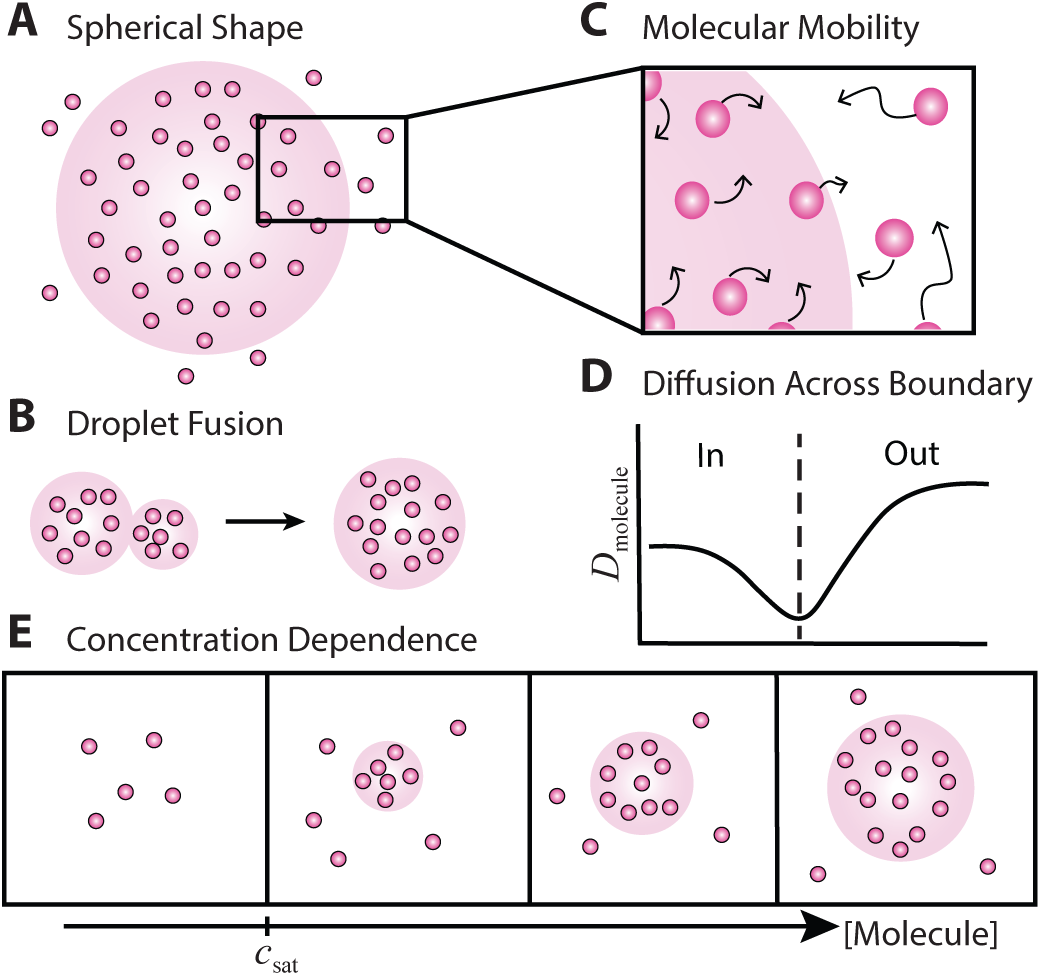
Proposed criteria to determine if a cluster assembles by LLPS. (a) Condensates are spherical due to surface tension. (b) Droplets can fuse upon contact. (c) Mobility of condensate components and their exchange across the boundary. Arrow length denotes the rate of diffusion of the molecule. (d) Restricted diffusion across the membraneless boundary. (e) Condensate formation and size scales with component concentration; cytoplasm component concentration is buffered.

Several bacterial protein-protein and nucleic acid-protein complexes have demonstrated liquid-like properties that may affect cellular function. For example, bacterial ribonucleoprotein bodies (BR-bodies) are condensates that contain the RNA degradosome. This condensate selectively sequesters mRNA to enhance RNA decay (17, 18). Another reported LLPS candidate is FtsZ, a key divisome assembly protein, which forms droplets in the presence of nucleoid occlusion factors *in vitro* (19). (19). FtsZ has been studied for decades in a number of bacterial species, yet condensate formation has thus far not been shown *in vivo*, leaving the role of LLPS in cell division and its spatial regulation a mystery. Further characterization of these and other hypothesized condensates will ascertain their biochemical functions and how they are regulated in time and space.

LLPS is recognized in eukaryotic cells when clearly visible spherical droplets demonstrate liquid-like properties through fusion or by external manipulation. However, these behaviors are almost impossible to observe in tiny bacteria using conventional microscopy. Thus, bacterial condensate formation has been demonstrated by *in vitro* reconstitution, by visualizing the building blocks with fluorescence or differential interference contrast microscopy, and by correlating these LLPS activities *in vitro* with the behaviors of foci in cells (17–26).

Additionally, the *in vitro* concentrations and environmental conditions (salt concentrations, pH, temperature, crowding agents) under which components phase separate have been measured (17, 19, 21–25, 27). The liquid-like behavior of condensates has been tested by fluorescence recovery after photobleaching (FRAP) and by observing droplet fusion events (17, 21, 24, 26, 28). Finally, 1,6-hexanediol, a compound known to dissolve condensates, has probed LLPS (23, 29), though hexanediol sensitivity may not suffice to demonstrate formation by LLPS (8). While these experiments support phase separation *in vitro* and indicate condensate formation in cells, they do not definitively show causality: a more rigorous diagnostic is necessary to specify that LLPS is responsible for cluster formation in bacteria.

Live-cell experiments are therefore needed. Super-resolution microscopy and related modalities have localized bacterial condensates *in vivo* (18, 21, 26, 30) and measured the diffusive properties of their molecular components (23, 26, 31, 32). As the methods improve and rigorous controls are implemented, single-molecule methods are becoming increasingly critical for understanding bacterial subcellular organization due to the very small size of the relevant features in these microscopic organisms.

Overall, the proposed criteria provide a framework with which to evaluate phase separation *in vivo*, particularly in bacteria. In this review, we describe the bacterial systems with proposed LLPS activity and discuss the experimental procedures used to investigate them. Furthermore, we comment on the general role LLPS plays in bacteria and how it may be part of a series of phase transitions that regulate cellular function. Lastly, we provide an outlook for super-resolution microscopy and single-molecule tracking as tools to assess condensates in bacteria.

### Current evidence demonstrates that LLPS may mediate subcellular organization in bacteria

#### mRNA degradation

To date, the best characterized prokaryotic condensates are the bacterial ribonucleoprotein bodies (BR-bodies), which control mRNA decay by locally concentrating the RNA degradosome proteins and their substrates (Fig 2A) (17, 18). In *Caulobacter crescentus*, assembly of this machinery is facilitated by RNase E which uses its C-terminal domain (CTD) as a scaffold (33). This essential domain for BR-body assembly is an IDR that contains many RNA-binding sites (17, 33). BR-bodies respond dynamically to decreases in translation levels by forming condensates that accelerate mRNA decay (18). Moreover, condensate disassembly is promoted by mRNA cleavage, which then releases mRNA fragments for other uses. For a more in-depth review of BR-body function and phylogenetic distribution, see Muthunayake et al. (4).

**Figure 2.**
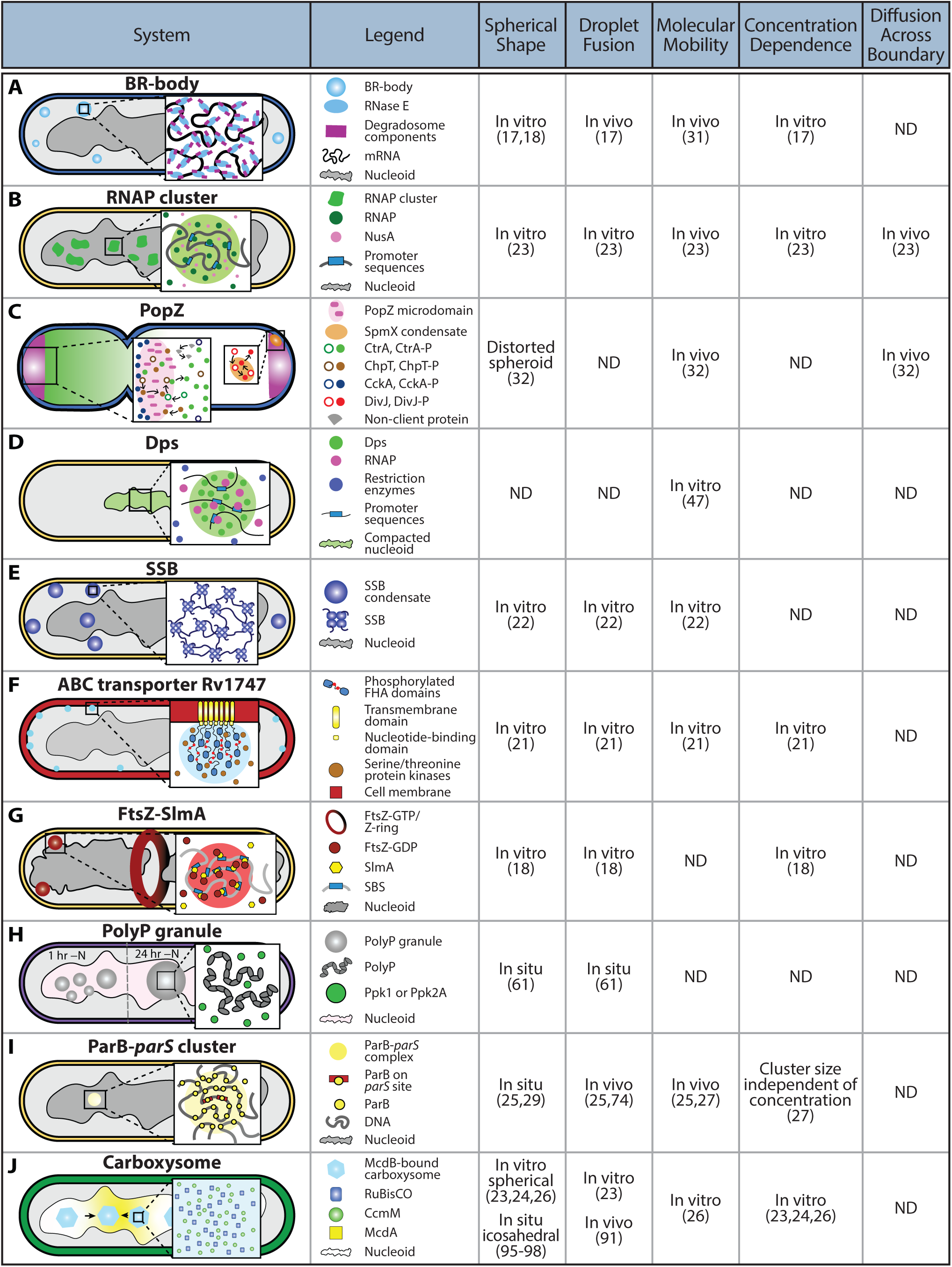
Proposed bacterial LLPS systems and evidence for the criteria in Figure 1. References are given in the figure. ND: not determined. Not shown: The α-carboxysome CcmM analog CsoS2 can phase separate *in vitro*. DEAD-box RNA helicases were also briefly discussed but not shown in the figure.

BR-bodies have liquid-like properties. RNAse E CTD, the essential component for BR-body assembly, forms spherical droplets *in vitro*, and RNA enhances RNAse E CTD droplet formation (17). Specifically, long, unstructured RNAs are preferentially recruited to BR-bodies and appear to mediate droplet size (18). *In vivo*, RNase E forms foci in different alpha-proteobacteria; these foci do not form under mRNA depletion (17). Moreover, RNase E CTD foci coalesce into larger spherical structures in *Escherichia coli*.

The molecular mobility within BR-bodies was characterized by single-molecule tracking of RNase E-eYFP in *C. crescentus* (31). Though many RNase E-eYFP copies are confined to the foci, most of these clustered RNase E-eYFP molecules exhibit confined motion with non-negligible diffusion coefficients. *In vitro*, spherical RNase E CTD droplets form only at NaCl concentrations lower than 200 μM, indicating a role for electrostatics in condensate formation. Also, the ability of BR-bodies to recruit certain RNAs and to exclude the ribosome and nucleoid support their selective permeability (18). This specific recognition may be a result of surface tension controlled by specific mRNA sequences or by mRNA concentration (34). How the diffusive properties of different RNAs or excluded ribosomal proteins change in or around the BR-body boundary has yet to be determined. Taken together, the evidence summarized here supports BR-bodies as bacterial condensates.

#### Transcription

RNA polymerase (RNAP), the enzyme responsible for RNA transcription, forms dense foci in fast-growing *E. coli* cells (35). This localization pattern has been linked to nucleoid organization (36). A recent study proposed that within these clusters, two types of RNAP exist: (1) actively transcribing RNAP and (2) non-transcribing RNAP (Ladouceur et al., 2020). Confining the non-transcribing RNAP to clusters maximizes ribosomal RNA (rRNA) transcription levels by accelerating re-initiation rates. Because they exhibit some of the hallmark features, Ladouceur and colleagues suggest that the RNAP clusters form via LLPS (Fig 2B). The dynamics of RpoC, an RNAP component, suggests a discrete transition from diffuse to clustered molecules, indicating a first-order phase transition (23). Moreover, RNAP clusters dissolve in the presence of 1,6-hexanediol, whereas overexpression-induced aggregates are unaffected by this perturbation, suggesting that RNAP clusters form via weak protein-protein interactions in addition to DNA binding. Notably, NusA, an antitermination factor that interacts with RNAP, forms droplets that fuse upon contact *in vitro* and nucleate foci independent of DNA binding *in vivo*. Lastly, single-molecule tracking revealed that RpoC and NusA remain mobile while confined to RNAP clusters and exhibit two diffusive rates, corresponding to the transcribing and non-transcribing populations. Collectively, these data support LLPS as a way to regulate nutrient-dependent transcription, though further investigations connecting this behavior to transcriptional pathways are necessary to uncover a complete functional picture.

#### Cell polarity

The asymmetric bacterium *C. crescentus* has two different cell poles: the new swarmer pole and the old stalked pole. Asymmetric activity of the phosphorylated transcription factor CtrA (CtrA-P) is required to produce this functional asymmetry in *C. crescentus*. Among the factors that contribute to generating this gradient, the polar organizing protein PopZ plays a large role. PopZ localization is bipolar after chromosome segregation and recruiting proteins to both old and new poles, but its effect is different at the two poles. At the stalked pole, PopZ forms a microdomain that is necessary for polar localization of a number of proteins, including various phospho-signaling proteins (37–39). A combination of fluorescence imaging and reaction-diffusion measurements demonstrated that the PopZ microdomain at the swarmer cell pole concentrates CtrA transcription factor signaling pathway proteins (Fig 2C). This high concentration enhances CckA kinase activity, increases the probability of intermolecular interactions to facilitate phosphate transfer by ChpT-P, and releases phosphorylated CtrA (CtrA-P) to generate a gradient (40, 32). Because the CtrA-P gradient decays away from the swarmer-cell pole, it results in a skewed inheritance of CtrA-P in the two daughter cells after asymmetric cell division (32, 40). This distribution ultimately leads to a differential expression of genes between the daughter cells and inhibition of replication initiation in the swarmer cell (41).

The evidence that supports this model for CtrA-P patterning, as well as the large disordered regions of PopZ, also suggests that PopZ microdomains are bacterial condensates (32). Super-resolution imaging of photoactivatable mCherry (PAmCherry), as well as correlative microscopy experiments (42) showed that PAmCherry-PopZ forms a dome at the cell pole while its cytoplasmic end is a membraneless interface. Single-molecule tracking found that the CtrA signaling pathway proteins have different diffusive behaviors on either side of the interface: CckA, ChpT, and CtrA diffuse more slowly within the swarmer-pole PopZ microdomain than in the cytoplasm. This difference in mobility results in longer dwell times in the swarmer pole microdomain, allowing for intermolecular binding and phosphate transfer; a conclusion further corroborated by simulations. Moreover, the microdomain is impermeable to free cytoplasmic proteins and requires proteins to associate with PopZ directly or indirectly for entry. Though the PopZ studies indicate certain key characteristics of condensates (Fig 1), further characterization of PopZ microdomains is required to converge on LLPS as the governing mechanism for their assembly.

For instance, the stalk and swarmer PopZ condensates act as an LLPS hub for different processes. At the stalked pole, an integral membrane protein, SpmX, interacts with PopZ and localizes the kinase DivJ, thus generating another asymmetry in the cell (43, 44). A recent study by Saurabh and colleagues used *in vitro* reconstitution in solution and on supported lipid bilayers to show that both PopZ and SpmX form condensates via LLPS (45). Furthermore, after heterologous expression of PopZ and SpmX in *E. coli*, it was found that diffusion of these proteins within the condensates in living cells could be modulated by changing any parameter that affects protein self-interaction (pH, osmolarity, crowding, etc.). Finally, the IDRs of PopZ and SpmX were found to be necessary and sufficient for the *in vitro* phase separation. Based on these experiments and on recent correlative cryogenic electron tomography (cryoET) images of SpmX and PopZ (42), it was proposed that PopZ and SpmX form two interacting membraneless organelles, one cytoplasmic (PopZ) and the other tethered to the membrane (SpmX) at the stalked pole (Fig 2C). Since SpmX phase separation slows DivJ diffusion to increase the polar concentration of DivJ (45), one of the functions of the SpmX condensate is to enhance the density-dependent DivJ kinase activity. It is therefore likely that two interacting membraneless organelles in *C. crescentus* robustly regulate the key DivJ signaling reaction.

#### DNA compaction

One of the most abundant proteins in the *E. coli* nucleoid, Dps (DNA-binding protein from starved cells) (46), is likely organized by phase separation (47). When Dps is highly expressed in stationary phase, it compacts the nucleoid to protect it from damage (48). However, DNA condensation by Dps does not affect transcriptional regulation and has negligible effect on the proteome in the stationary phase (47). However, selective accessibility to DNA was observed upon Dps-induced compaction *in vitro*. While RNAP maintains its ability to bind its promoters, DNA saturation with Dps blocks the activity of certain restriction enzymes. It is therefore hypothesized that Dps achieves this selectivity by forming a phase-separated DNA-Dps complex: unlike other enzymes, RNAP enters and diffuses freely within these complexes, which suggest selective permeability (Fig 2D) (6, 18, 47). Additionally, *in vitro* single-molecule assays demonstrated rapid rearrangement of Dps complexes. The Dps mobility suggests that the phase of the Dps-DNA complex may be tunable and responsive to condition changes that promote liquid-like behaviors.

#### DNA repair

Single-stranded DNA binding protein (SSB) stabilizes single-stranded DNA (ssDNA) and recruits proteins essential for DNA replication, repair, and recombination (49). This NAP is a good candidate for condensate formation through LLPS because it forms phase-separated droplets, is promiscuous in its binding interactions, and contains an IDR (22). A combination of turbidity measurements and fluorescence imaging found that SSB forms liquid-like droplets *in vitro*. Notably, SSB droplets can fuse and are apparently viscous based on FRAP measurements. Moreover, droplet size appears to depend on SSB concentration up to a certain threshold concentration, with or without crowding agents. Further genetic and fluorescence studies determined that the C-terminal domain of SSB is crucial for phase separation due to its interaction with the OB fold, and that this interaction is inhibited by saturation with ssDNA (22). Harami and colleagues proposed that as the amount of exposed ssDNA increases, membrane-localized SSB condensates (50) rapidly dissolve to localize SSB to these sites of increased ssDNA (Fig 2E). This transition may occur in response to genomic stress or upon initiation of DNA repair (22). If the critical role of SSB in DNA repair and its apparent ability to phase separate is validated, this phase separation is potentially a conserved mechanism in bacteria. Because SSB droplet formation is governed by weak protein-protein interactions, ssDNA is not crucial for SSB to recruit related proteins, allowing for membrane-localized condensates.

#### Transmembrane transporters

The ATP-binding cassette (ABC) transporter Rv1747 is a virulence factor important for *Mycobacterium tuberculosis* growth in infected hosts (51). Rv1747 contains FHA regulatory modules, which mediate its virulent activity, and phospho-acceptor threonines in the intrinsically disordered linkers (52). The FHA domains (Rv1747^1-310^) phase separate at high concentrations. Their phase separation is reversed by the addition of the *M. tuberculosis* phosphatase, PstP, and is induced by other related kinases. For instance, Rv1747^1-310^ phosphorylation by the PknF kinase enhances condensate formation *in vitro* (21). Moreover, a FRAP measurement of diffusive exchange of Rv1747^1-310^ condensates estimated a 60% recovery for both unphosphorylated and phosphorylated samples, but found that the recovery is much slower after phosphorylation. These data suggest that Rv1747^1-310^ diffusion in the condensate is limited by phosphorylation, possibly because this reaction enhances electrostatic interactions (53), which would also explain the induction of Rv1747 phase separation by PknF. Consistent with condensate formation, the coalescence of fluorescently tagged Rv1747^1-310^ into dense foci was also observed when Rv1747 was expressed in *S. cerevisiae* and *Mycobacterium smegmatis* (21).

Another potential connection between kinase and phosphatase cofactors and Rv1747 phase separation is their different localization patterns within the condensates *in vitro* (21). For example, PknF is distributed throughout the entire focus while PstP forms discrete puncta at the interface between the condensate and the surrounding solution. The discrete localization of PstP to the boundary suggests that it is unable to penetrate the condensate, implying a selectively permeable boundary that prevents phosphorylated Rv1747 condensates from dissolving, though the diffusive properties of PstP have not been measured. In contrast, permeability of the condensate to PknF further supports the role of PknF as a LLPS enhancer (53, 54). Finally, super-resolution images depicted Rv1747 nanoclusters at the membrane of *M. tuberculosis*. Taking this membrane clustering together with the phosphatase/kinase results led the authors to a working model in which these enzymes promote Rv1747 clustering at the membrane to increase transport efficiency, provide selective signals based on substrate permeability, and/or form scaffolds for interactions with cell wall biosynthesis components (Fig 2F) (21). Currently, it is unclear if phase separation drives the nanoclustering and it is possible that other driving mechanisms exist, but the prospect of LLPS playing a role in cell membrane functions is nonetheless exciting.

#### Cell division

FtsZ is a highly conserved and central player in the assembly of the bacterial cell division machinery (the divisome). This tubulin-like GTPase polymerizes into a Z-ring structure, which initiates cell division at the bacterial midcell (55). The Min system and other nucleoid occlusion factors like SlmA specifically position FtsZ to constrain division to this site (55, 56). Monterroso and co-workers proposed that prior to Z-ring formation, binding of SlmA to SlmA-binding sequences (SBS) helps sequester FtsZ within condensates near the cell membrane, and that once the cell is ready for division, FtsZ is recruited to the midcell to polymerize into the Z-ring driven by GTP (Fig 2G) (19).

Under crowded *in vitro* conditions with or without GDP, *E. coli* FtsZ forms droplets, and in the presence of both the polymerization antagonist SlmA and oligonucleotides containing SBS (19). To determine if FtsZ from the FtsZ-SlmA-SBS condensates could polymerize, GTP was added to FtsZ-SlmA-SBS condensates. Minutes after GTP addition, confocal microscopy indicated FtsZ polymerization with subsequent droplet disruption, likely due to soluble FtsZ binding GTP to form polymer seeds that decrease the solute concentration below *c*_sat_. Subsequent GTP hydrolysis rescues the condensate. Further experiments using microfluidic-generated microdroplets, which simulate cellular confinement, demonstrated that FtsZ condensates prefer the lipid boundary while FtsZ fibers localize to DNA-rich regions. This *in vitro* finding suggests that in live cells, FtsZ condensates may be excluded from the nucleoid. Taken together with previous work showing that FtsZ enhances assembly across crowded, phase-separated microenvironments (57), it is possible that these condensates act as FtsZ sinks to negatively regulate Z-ring formation; however, there is not yet evidence of FtsZ condensates *in vivo*. The intrinsically disordered C-terminal tail of FtsZ, which binds SlmA (56), has been proposed to function under the stickers-and-spacers framework to explain the multivalent interactions that drive phase transitions (58).

#### Cell starvation response

Another candidate for phase separation in bacteria is polyphosphate (polyP) granules. PolyP is an inorganic polymer consisting of phosphoryl groups; polyP granules are present in bacteria under stress and starvation (59, 60). For bacteria to exit the cell cycle to survive under stress, they must prioritize completing DNA replication (61). Due to its connection to the starvation-signaling molecule guanosine tetraphosphate, Racki and co-workers studied *de novo* polyP granule formation in *Pseudomonas aeruginosa* to determine if polyP protects the nucleoid under stress conditions (62). Following nitrogen starvation, polyP deletion leads to *P. aeruginosa* cell elongation, indicating a link between polyP and cell cycle exit during starvation. Moreover, the number of polyP granules decreases over time after nitrogen starvation, but the size of each granule increases at the same time, suggesting that the granules can fuse (Fig 2H). Interestingly, there also appeared to be a minimum distance between granules and the cell poles. This organization was investigated by fluorescence microscopy: Ppk2A-mCherry, a polyphosphate kinase responsible for polyP metabolism (63), and GFP-ParB, to mark the origin of replication, were visualized to determine if the nucleoid prevents granules from positioning at the cell poles (Fig 2H). Though the results did not determine if polyP granules perfectly colocalize with the chromosome origins, they did suggest that the nucleoid forces the granules to maintain a minimum distance from the cell poles.

The newly characterized role of polyP granules in cell cycle exit raises some questions about the specific biochemical function of these granules. Racki and colleagues proposed that these granules may serve as microcompartments that compartmentalize specific enzymatic activity, akin to the roles of condensates such as P bodies and nucleoli (6, 8). This connection is especially interesting considering the similarities in physical properties between polyP and the nucleic acid oligomers that are typically found in phase-separated bodies: both have high conformational flexibility and repetitive motifs capable of forming weak interactions with itself and other biomolecules (6, 9, 62). These physical parallels, in combination with the evidence suggesting that polyP granules fuse and may be positioned through some mechanism involving the nucleoid, imply that polyP may be a condensate that uses phase transitions to regulate its biochemical function.

## Condensate formation governs partition complex formation

The ParABS partition system is responsible for the segregation and faithful inheritance of most chromosomes and plasmids in bacteria (Fig 2I). ParB proteins form dimers that recognize and bind to centromere-like sites on DNA known as *parS* sites. This initial site-specific association with *parS* nucleates the recruitment of hundreds of ParB dimers that load onto and around the *parS* site through ParB-ParB interactions as well as through non-specific binding to DNA flanking the *parS* site (28, 30, 64–66). After the ParB-*parS* complex forms, replicated DNA is segregated by interactions between ParB and its corresponding ParA protein that coats the nucleoid via non-specific DNA binding. The large ParB-*parS* complex stimulates ParA ATPase activity, which is coupled to its local release from the nucleoid. The resulting ParA protein gradient directs the motion of DNA cargoes toward higher concentrations of ParA on the nucleoid via weak, transient interactions between ParB and ParA. Diffusion-based models have been proposed to explain the segregation, directional movement, and positioning of ParB-bound DNA cargoes via dynamic ParA gradients on the nucleoid (67–70).

How a small *parS* site, which allows for the site-specific binding of only a few ParB dimers, nucleates the loading of hundreds of ParB dimers to form a massive nucleoprotein remains unclear. Four models have been proposed: (1) In the spreading model, a ParB dimer specifically binds to *parS* and further ParB dimers interact with neighboring dimers to propagate in 1D along the DNA track (64); (2) In the spreading, bridging, and looping model, 1D spreading is expanded to include non-specific DNA bridging via ParB dimer-dimer interactions and subsequent DNA looping (65, 66); (3) In the nucleation and caging model, which arose from chromatin immunoprecipitation (ChIP) assays and mathematical modeling, ParB specifically binds to *parS* and a network of transient, weak interactions of ParB dimers with other dimers and with nsDNA is nucleated at *parS*. DNA near *parS* then enters this region of high ParB density, leading to stochastic binding of ParB that is consistent with the experimentally observed power law decrease of ParB concentration unveiled in the ChIP experiments (28, 30); (4) In the final model, based on recent work demonstrating ParB CTPase activity (71–73), ParB dimers form CTP-driven clamps after nucleation at *parS* via a conformational change, then ParB dimer clamps spread along flanking DNA via sliding (74).

Many pieces of evidence point to the dynamic ParB-*parS* complexes as condensates. For example, single-molecule localization microscopy (SMLM) experiments in combination with single-particle reconstruction demonstrated that the majority of ParB-mEos2 molecules localize to two dense, spherical clusters on plasmids located at the *E. coli* cell quarter positions (Fig 3A) (26, 30). Furthermore, ParB clusters are liquid-like in their ability to fuse, as demonstrated for the F and P1 plasmid partition systems in *E. coli* (26, 75). Interestingly, real-time imaging after ParA degradation revealed ParB cluster fusion, suggesting that when ParA interacts with ParB-bound condensates to distribute DNA cargoes, ParA prevents condensate coalescence (Fig 3B). The differential mobility of ParB inside and outside of the cluster further supports phase separation. Single-molecule tracking of ParB revealed a small (5 %) freely diffusive population and a large static population which spatially correlates to ParB-*parS* complex sites (Fig 3C) (26). Additionally, a combination of FRAP and fluorescence lifetime imaging microscopy measurements determined that the dynamic exchange between two ParB clusters occurs on the scale of a few minutes (26, 28). These data demonstrate that while ParB is primarily confined to the vicinity of a *parS* site, it maintains the ability to freely diffuse and even exchange to another ParB cluster, further supporting that this system is a bacterial condensate.

**Figure 3.**
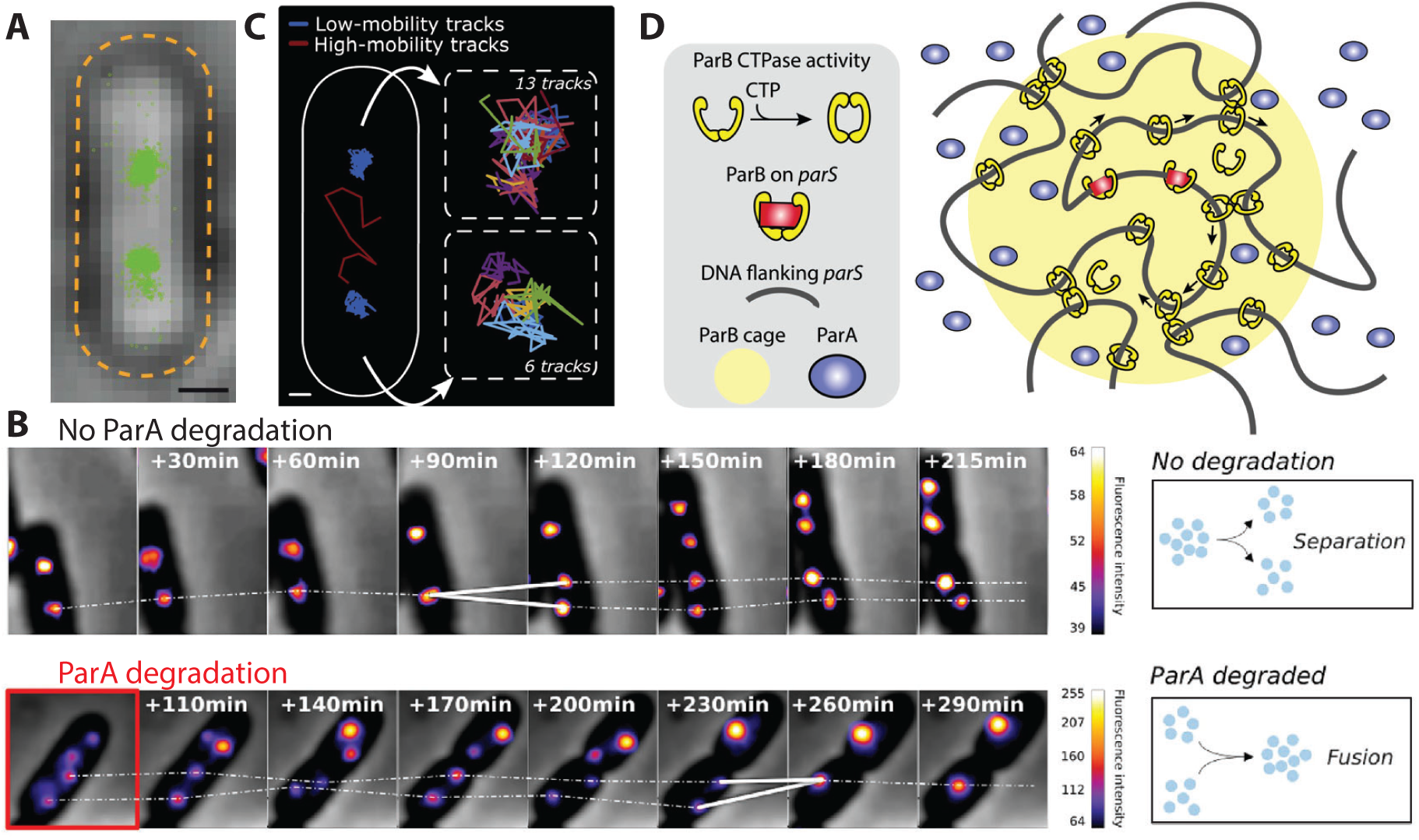
ParB forms clusters at *parS* sites, which are segregated by ParA. (a) Super-resolution images of ParB clusters. Scale Bar: 500 nm. From ref. (30). (b) In cells expressing ParB-mEos2 and ParA fused to the SsrA degron tag, time-lapse images of a ParB cluster separating without induction of ParA degradation (top) and fusing upon induction of ParA-Ssra degradation (bottom). From ref. (26). (c) Representative ParB single-molecule tracks From ref. (26). (d) A hybrid model for ParB condensation at *parS* that includes aspects from the ParB nucleation and caging model (refs. (28, 30)), ParB CTPase activity (ref. (73)), and potential interplay between ParB CTPase and ParA ATPase activities.

Beyond the correlative evidence supporting ParB-*parS* complexes as condensates, phase separation can explain related functional roles. For example, weak interactions between ParB and ParA drive ParA-gradient asymmetry by stimulating its ATPase activity and release from the nucleoid (67, 69, 70, 74, 76–78). How a plurality of weak interactions manages to create enough mechanical force to segregate newly replicated chromosomes or plasmids remains unclear. In the diffusion-ratchet model, the mechanochemical pulling force is provided by hundreds of ParA-ParB interactions at the surface of the ParB-*parS* complex (79, 80). It is unknown whether ParA can permeate into this complex, which would allow for an even greater interaction density with ParB dimers. This partition process may operate near a critical point, which would allow the system to detect and respond to changes in nucleoid size and plasmid copy number (81). Additionally, partition complexes may buffer local ParA concentrations to maintain partition fidelity upon variations in cellular ParA levels (81); this buffering is one LLPS indicator (15).

An alternative hypothesis to partition-complex formation by LLPS is assembly by polymer-polymer phase separation (PPPS) (82). In this model, the partition complex size is governed by the DNA length rather than by the ParB concentration (83). Accordingly, modeling data suggest that partition complex size is independent of the ParB concentration (28). Further evidence for PPPS is that in *Bacillus subtilis* and *Streptococcus pneumoniae*, ParB helps to recruit structural maintenance of chromosome (SMC) complexes (84–86), which compact and resolve replicated chromosomes (87). *Saccharomyces cerevisiae* SMC has recently been shown to form bridging-induced phase separated bodies akin to ParB PPPS (88). Finally, given that *in vitro* reconstitution of ParB condensation on *parS* substrates has been unsuccessful, it is likely that CTP is the missing piece of the puzzle. While the ParB CTP-driven spreading model is convincing, we propose that these models are not mutually exclusive. Rather, this CTP-binding and CTPase activity of ParB provides an energy source as well as a molecular view of the nucleation and caging model, wherein the spreading that leads to the caging of *parS* by ParB dimer-dimer interactions is explained by ParB sliding (Fig 3D). We also speculate that ParA may influence ParB CTPase activity such that ParA concentration regulates partition complex size in addition to its positioning in the cell.

## LLPS may be involved in carboxysome biogenesis and positioning

Many bacteria sequester chemical reactions or store important molecules through protein-based organelles known as bacterial microcompartments (BMCs) (89). Rather than using a lipid membrane, a selectively permeable protein shell forms the boundary between the BMC contents and the cytosol (Fig 2J). Carboxysomes, the carbon-fixing BMCs found in photosynthetic cyanobacteria as well as many chemoautotrophic bacteria, are responsible for ∼35% of global carbon fixation (90). Carboxysomes efficiently fix carbon by encapsulating ribulose-1,5-biphosphate carboxylase/oxygenase (RuBisCO), the enzyme that catalyzes the first major step of the Calvin cycle. The carboxysome protein shell is permeable to HCO_3_^−^, which is converted into CO_2_ by carbonic anhydrase (CA) also encapsulated in the carboxysome (91). This mechanism is more formally known in cyanobacteria as the Carbon Concentrating Mechanism (ccm) (91).

In the last decade, fluorescence microscopy has measured carboxysome assembly and organization (92–95) to complement the extensive structural characterization of BMCs (96–98). Though carboxysomes were previously believed to be exclusively encapsulated by a paracrystalline protein shell, recent clues suggest that LLPS may play a role in both carboxysome biogenesis and regulation (24, 25, 27).

### Carboxysome biogenesis and composition

The major players in carboxysome formation *in vivo* can form droplets *in vitro* (25, 27). The short form carboxysome protein M35 can form LLPS droplets with RuBisCO *in vitro* (Fig 4A) (27). FRAP measurements of both reduced and oxidized M35 droplets found recovery on the scale of a couple of minutes, though the oxidized M35 recovered more fully and twice as fast as the reduced M35. These results suggest that oxidized M35-RuBisCO is more dynamic than its reduced counterpart, which coincides with the proposed oxidized state of mature carboxysomes (93). Moreover, cryoET finds that RuBisCO-M35 distributes in dense clusters that closely resemble the liquid-like RuBisCO-EPYC1 condensates of the algal pyrenoid (99, 100), which is the functional eukaryotic analog of the carboxysome.

**Figure 4.**
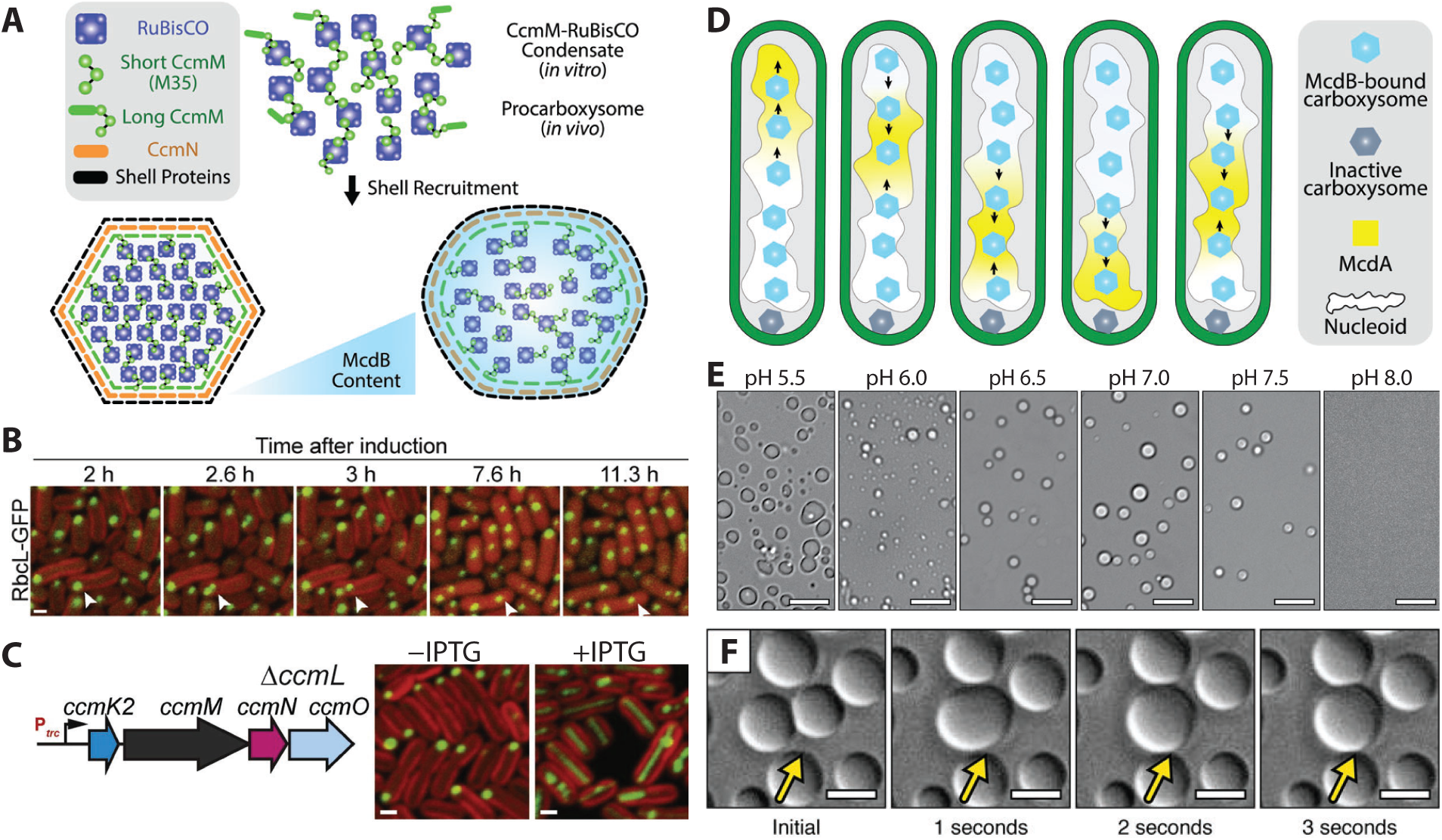
LLPS may play a role in carboxysome biogenesis and organization. (a) Model for CcmM-RuBisCO nucleation. Once assembled, carboxysomes may be fluidized by McdB. (b) Progression of carboxysome formation following induction of the *ccm* operon in *ΔccmK2-ccmO S. elongatus* expressing RbcL-GFP (green). White arrow denotes budding event from a procarboxysome. Scale bar: 1 μm. From ref. (92). (c) RbcL-GFP (green) forms bar carboxysomes upon CcmL deletion (+IPTG). Scale bars: 1 μm. From ref. (92). (d) Carboxysome positioning is governed by the McdAB system via a Brownian ratchet. Inactive carboxysomes, possibly with no McdB, have been shown to become polarly localized. (e) Microscopy images of *S. elongatus* McdB droplets under varying pH. Scale bars: 10 μm. From ref. (24). (f) McdB droplet fusion events (yellow arrows). Scale bars: 5 μm. From ref. (24).

There are two types of carboxysome: α and β. *S. elongatus* PCC7942 is a model cyanobacterium for the study of β-carboxysomes. Cameron and colleagues monitored carboxysome biogenesis by time-lapse fluorescence microscopy and observed that the large subunit of RuBisCO (RbcL) coalesces into aggregates following *ccm* operon induction. The shell is formed after core components assemble into a “procarboxysome” (Fig 4A) and before CcmL is incorporated; ultimately the mature carboxysome is released from the procarboxysome. Interestingly, this proposed birth of new carboxysomes from the procarboxysome comes from observations of budding and of the linear relationship between time and carboxysome copy number (Fig 4B) (92, 93). Furthermore, CcmL is necessary for carboxysome budding and its deletion leads to elongated “bar” carboxysomes (Fig 4C). Less is known about α-carboxysome assembly, however the ability of their shell components to form empty carboxysome “ghosts” suggests that the inside-out assembly pathway of β-carboxysomes is not obligatory for α-carboxysome assembly (101, 102). Still, the N-terminal domain of CsoS2, a required protein for α-carboxysome assembly, can form phase-separated droplets with RuBisCO (25). Since CsoS2 is an intrinsically disordered protein (IDP) that is believed to act as a scaffold for RuBisCO coalescence (103), it is possible that like CcmM, CsoS2 might use LLPS to initiate carboxysome formation.

Carboxysome structure, composition, function, and organization are all highly responsive and adaptable to environmental change including changes in growth temperature, light, and CO_2_ concentration (94, 104, 105). For example, under high light intensities, the number of *S. elongatus* carboxysomes doubles from ∼5 to 10, whereas the number decreases to one or two per cell under low light (94, 106). Indeed, irradiance increases carboxysome size and component production, and consequently increases levels of carbon fixation. Furthermore, single-molecule slimfield microscopy detected that the abundance of carboxysome proteins changes as a function of light and CO_2_ levels (105).

### Carboxysome positioning

The rod-shaped *S. elongatus* evenly distributes carboxysomes along its long axis (106). <u>M</u>aintenance of <u>c</u>arboxysome <u>d</u>istribution protein <u>A</u> (McdA) is a ParA-type ATPase that establishes the oscillatory gradients responsible for carboxysome organization in the cell when its partner McdB interacts with carboxysomes and prompts McdA ATPase activity and release from the nucleoid (95, 106). This McdA displacement creates an asymmetric distribution of McdA proteins, and carboxysomes are driven toward regions of higher McdA concentration (Fig 4D) (95). The carboxysome motion is therefore directed by a Brownian ratchet mechanism akin to ParA-based DNA segregation systems (69, 95). Moreover, the McdAB system is widespread among β-cyanobacteria and carboxysome-containing proteobacteria, suggesting that carboxysome positioning is a conserved process across these groups (24, 107).

Deletion of *mcdA* or *mcdB* leads to a wider distribution of carboxysomes in the axial direction and larger mean focus size due to carboxysome aggregation (95). The data also suggest that without the positioning system, newly formed carboxysomes cannot be separated and readily fuse with other carboxysomes (92, 95). McdB overexpression results in the formation of bar carboxysomes similar to those reported in cells lacking CcmL (92), and may be the result of an inability to bud new carboxysomes. Bacterial two-hybrid assays suggest that an ensemble of interactions govern the role of McdB in carboxysome organization (95), but how McdB interacts with carboxysomes remains unclear.

McdB can undergo LLPS *in vitro* and this condensation has been proposed to be involved in carboxysome organization (Fig 4A) (24). McdB proteins across β-cyanobacteria share common features, such as IDRs, repetitive and biased amino acid sequences, low hydrophobicity, and high multivalency with proteins that undergo LLPS (6, 24). *S. elongatus* McdB can form pH-dependent liquid-like droplets *in vitro* (Fig 4E) (24) and neighboring McdB droplets fuse (Fig 4F). At lower pH, McdB droplets are gel-like, and at higher pH, McdB does not undergo LLPS. It has been proposed that the metabolic activity in the carboxysome lumen correlates with a carboxysome local environment that is more acidic than the cytosol (108, 109).

Therefore, it is attractive to speculate that carboxysome pH not only allows for McdB recruitment via LLPS, but also allows McdB to be selectively recruited to functionally active carboxysomes: carboxysomes that are metabolically inert would not have the low pH required for McdB recruitment and would no longer be positioned by McdA (Fig 4D). Consistent with this proposal, Hill et al. recently found that inactivated Synechococcus sp. PCC 7002 carboxysomes are mispositioned toward the cell poles and rapidly degraded (110). Cyanobacteria may use McdAB positioning systems to sense which carboxysomes are still active and require positioning and which should be targeted for degradation. Determining if McdB indeed forms condensates *in vivo* and what functions it serves beyond carboxysome positioning is an exciting future direction.

### Carboxysomes as bacterial condensates

Carboxysome LLPS has not been reported *in vivo*, but current cytological evidence certainly suggests that carboxysomes are bacterial condensates. *In vivo*, RuBisCO and CcmM nucleate into dense foci that resemble known condensates. However, the spatial resolution of standard fluorescence imaging is insufficient to determine if these foci are indeed spherical in shape (92, 93). Still, microscopy has observed pairs of adjacent foci that interact such that one is depleted as the other increases in size (92). In combination with the splitting events that demarcate carboxysome formation (Fig 4B), these observations are consistent with liquid-like behavior, and the spatial organization of such condensates may be regulated by the McdAB system (Fig 4D) (24, 95). Furthermore, carboxysomes can respond to their environment: their composition and dynamics are modulated in response to the photosynthetic activity of cyanobacteria (105). It is still unknown if and how LLPS plays a role in carboxysome biogenesis, organization, and regulation of carbon-fixation.

## Phase transitions enable adaptation to subtle changes in the cellular environment

Increasing evidence in eukaryotes and prokaryotes indicates that phase transitions can provide adaptive response to small changes in the cellular environment. For example, inducing phase separation in response to changes in pH or salt concentration may control protein expression (111). In particular, the bacterial cytoplasm can restrict component mobility in a size-dependent manner by exhibiting glass-like properties during lower metabolic activity (112). On the other hand, cellular metabolism fluidizes the cytoplasm and increases component motion, an effect more evident for larger particles. It will be interesting to consider the interplay between phase transitions in the cytoplasm and phase transitions of the membraneless organelles within.

For example, ATP depletion may lead BR-bodies to transition to a gel or solid state through decreased ATP hydrolysis by DEAD-box RNA helicases (4). Among these helicases are RhlB, a major degradosome component (33), and RhlE, whose degradosome association is crucial for fitness when *C. crescentus* is grown in cold temperatures (113). Furthermore, *E. coli* RhlE-mCherry forms condensates, whereas RhlB-mCherry lacks this ability (20). Perhaps stress or reduced metabolism causes BR-body phase transition to store mRNA, akin to p-granules or stress granules in eukaryotes (114, 115).

Carboxysomes may also take advantage of phase transitions. The increased carboxysome motion upon higher exposure to light may be related to the increased metabolic activity that follows light exposure in cyanobacteria (94, 105): the fluidized cytoplasm may encourage carboxysome motion. Furthermore, a fluid cytoplasm may drive interactions between carboxysome components, such as RuBisCO and CcmM, to increase nucleation and subsequent carboxysome formation, while a glassy state would hinder these interactions (92, 112, 116). Historically considered as paracrystalline structures with a protein shell (91), carboxysomes are beginning to be considered as more liquid-like (24, 25, 27). We posit that these phases are not mutually exclusive, but instead a spectrum of material states can exist depending on the cellular environment. It is possible that during metabolic dormancy, carboxysomes adopt their paracrystalline state to limit reaction rates (111). Indeed, the faceted structures of the observed icosahedral carboxysomes may be a low-energy arrangement, akin to that shown for elastic vesicles (117). In contrast, converting to a fluid composition would enhance reaction rates in carboxysomes and control the transport of substrates into and out of the protein shell. It is attractive to speculate that this shift in view is enabled by the methodologies used to study these compartments. Pioneering carboxysome imaging by transmission electron microscopy (TEM) required cells to be fixed, stained, and dehydrated (96). On the contrary, both fluorescence and cryoET microscopy indicate that carboxysomes are more structurally dynamic (99, 105), thus demonstrating the need for *in vivo* and *in situ* methods to definitively probe the behavior of these hypothesized condensates.

## Revisiting the assembly and material properties of bacterial inclusion bodies in the era of LLPS

Bacterial inclusion bodies (IBs) are mesoscale protein aggregates that are primarily formed by recombinant protein. Largely considered a bottleneck for producing soluble protein, detailed studies of IBs have been historically hindered by their presumed status as an overexpression artifact and by purification hurdles. Several strategies ranging from harsh denaturation to mild extraction have been developed to retrieve functional protein from IBs (118). IBs were once thought to be formed exclusively by misfolded or unfolded proteins, which aggregate into non-functional and nucleoid-excluded protein clusters (119). After 2005 however, dynamic IB models emerged in which amyloidal proteins form the IB and then offer mechanical stability to entrap non-amyloidal functional protein (120).

LLPS has only recently been considered as a way to understand the intriguing molecular organization and assembly pathway of IBs. For instance, the measles virus was recently shown to form inclusion bodies with properties of liquid organelles (121). In addition, numerous proteinaceous condensates have been shown to mature into semi-reversible gels and irreversible amyloids (1). Therefore, in addition to direct aggregation mechanisms, LLPS may be involved in forming bacterial IBs. However, direct determination of whether a massive focus is a liquid, gel, solid, or a mixture of these states is difficult in bacteria. One issue is that measurements used *in vitro* to show that a focus has liquid-like properties (Fig 1), cannot generally distinguish between liquid and solid aggregates in bacteria. For instance: droplets and solid bacterial aggregates are both spherical; the size of droplets and aggregates both scale with protein concentration; an increasing fluorescence signal may indicate droplets undergoing FRAP recovery, but this signal can also be attributed to a growing aggregate; and droplets fuse, but aggregates can also snap together. To differentiate between these phase states *in vivo*, it is imperative to resolve focus features that are exclusive to liquids: boundary effects on diffusion (Fig 1C-D), a buffered soluble phase, reversibility in response to changes in growth conditions (i.e., temperature shifts), and focus deformation in response to shearing forces. Due to the small size of these features, as we detail below, we believe super-resolution fluorescence imaging is critical for assessing LLPS in bacterial cells.

## Super-resolution microscopy as a tool to assess LLPS in bacteria

To date, most investigations of LLPS in bacteria have relied on correlative and qualitative observations. The typical workflow consists of identifying proteins with IDRs within diffraction-limited foci in bacteria, purifying these components to examine their liquid-like properties such as droplet fusion, and using FRAP to demonstrate dynamic molecular motion (8, 122). Unfortunately, the diffraction limit of light blurs images to ∼1 μm resolution, making nearly all subcellular droplets in bacteria appear spherical (122). Moreover, FRAP cannot ascribe mechanism. For instance, this method cannot tell if the cause of observed fluorescence recovery is free diffusion back to the photobleached position or specific binding at that site (122).

Therefore, just as our overall current understanding of bacterial cell biology would not have been complete without the emergence of super-resolution microscopy (123, 124), so too is super-resolution fluorescence imaging instrumental for assessing LLPS in bacterial cells (Fig 5). In particular, single-molecule localization microscopy (SMLM) methods, such as <u>P</u>hoto<u>a</u>ctivated <u>L</u>ocalization <u>M</u>icroscopy (PALM) and <u>St</u>ochastic <u>O</u>ptical <u>R</u>econstruction <u>M</u>icroscopy (STORM), are powerful because they determine how proteins behave in time and space by localizing single emitters with 10 – 30 nm localization precision and 10 – 100 ms time resolution; recent advances have pushed this limit down to a single nanometer (125, 126). Thus, super-resolution imaging and single-molecule tracking are ideal techniques to probe the localization and dynamics of LLPS in small bacterial cells.

**Figure 5.**
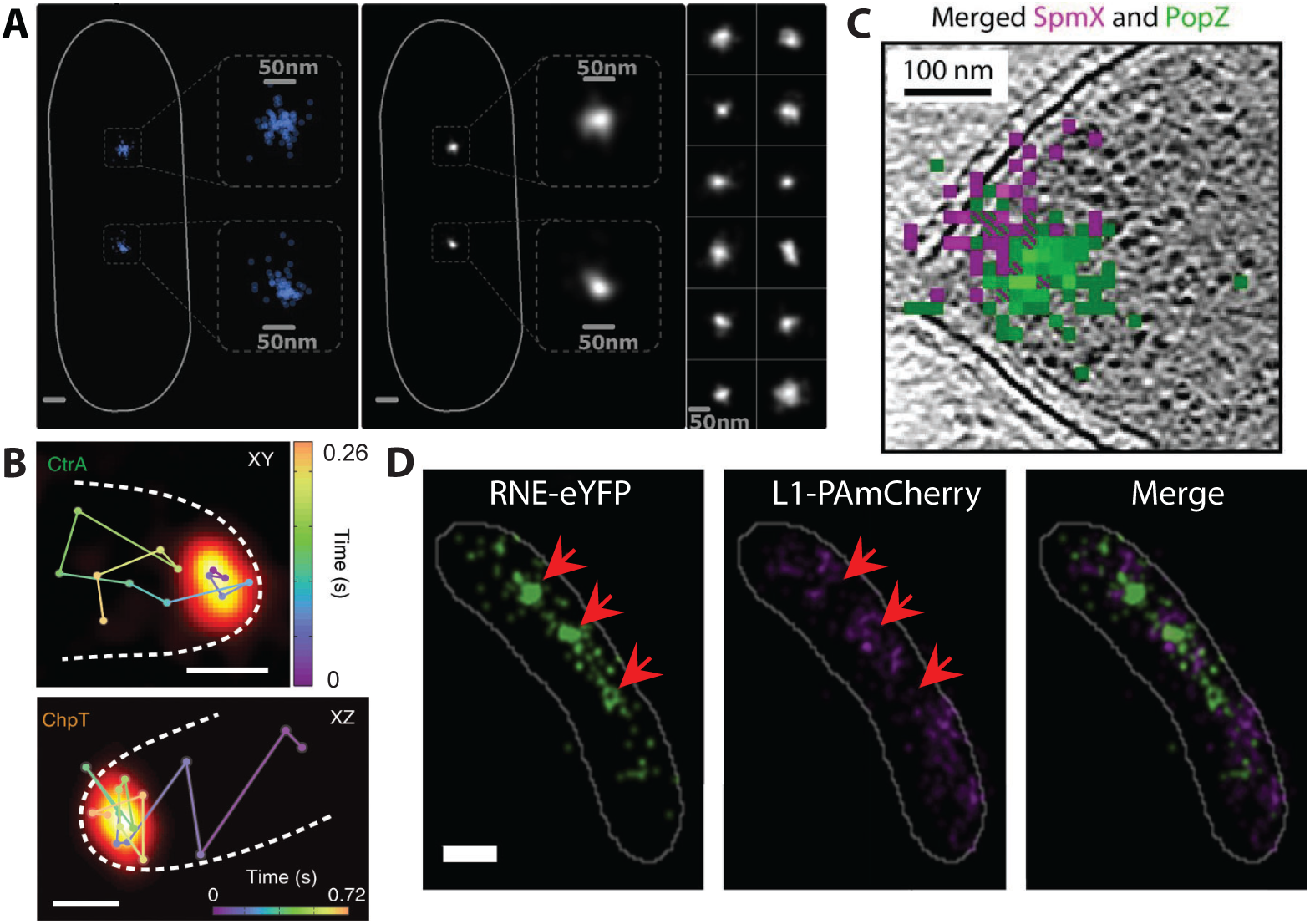
Single-molecule tracking characterizations of molecular condensates *in vivo*. (a) Super-resolution reconstructions of ParB clusters in *E. coli* (left) are class-averaged to estimate size, shape, and concentration (right). From ref. (26). (b) Super-resolution localizations of SpmX (violet) and PopZ (green) are overlaid with an electron tomogram of the *C. crescentus* cell at the stalked pole. From ref. (42). (c) 3D single-molecule tracks (time-coded connected dots) overlaid with the super-resolution images of PopZ (orange) in *C. crescentus*. Scale bars: 200 nm. From ref. (32). (d) Super-resolution images of RNaseE-eYFP (green) and ribosomal component L1-PAmCherry (violet) in *E. coli*. Arrows denote regions of ribosome exclusion with BR-bodies. Scale bar: 500 nm. From ref. (18).

In eukaryotes, super-resolution imaging and single-molecule tracking revealed that viral replication compartments are not assembled through LLPS since they form at different concentration levels, internal constituents are not evenly or randomly distributed, and their rate of diffusion inside of the compartment is equivalent to the diffusion rate in the nucleoplasm (127). Accordingly, examinations of protein clusters in bacteria have been applying these methods to quantitatively assess LLPS (23, 26, 31, 32, 128). We propose that these implementations need to be expanded and broadly applied to rigorously assess the previously described criteria (Fig 1) to confirm or rule out LLPS as a principal driver of spatial organization in bacteria.

To confirm that a condensate is spherical, the fine structure of proposed bacterial condensates can be determined with super-resolution microscopy. To further enhance the structural resolution, single-particle reconstruction algorithms, traditionally used in cryogenic electron microscopy, have been implemented (129). For example, this technique was used to estimate the shape, size, protein copy number, and concentration of ParB clusters (Fig 5A) (26). Advances to this computational analysis now permit single-particle reconstruction from multicolor SMLM data (130), which is crucial for understanding multi-protein systems like RNAP clusters and carboxysomes. It should be noted that further super-resolution methods development is needed—for instance, SMLM for the study of carboxysomes in cyanobacteria is challenging because of the high background fluorescence that results from its photosynthetic pigments (131).

Cryogenic super-resolution imaging is another promising technique that acquires structural details at a molecular resolution. By using super-resolution fluorescence imaging to identify target proteins in cryoET reconstructions, cryogenic super-resolution imaging has already revealed the precise localization of the PopZ-SpmX signaling complex (Fig 2C) within *C. crescentus* polar condensates (Fig 5B) and this method promises to determine extremely fine structural features and to measure the positioning of protein complexes in their native state (42).

Given that condensate constituents maintain their ability to diffuse inside the compartment, single-molecule tracking provides a quantitative alternative to FRAP measurements. This method has already been implemented in live *C. crescentus* and *E. coli* to investigate PopZ microdomains (Fig 5C) and RNAP clusters, respectively (23, 32). These studies revealed differences between the mobility of molecules inside and outside these compartments. Additionally, PopZ microdomains exclude cytoplasmic proteins, indicating an energetic barrier for entry. Super-resolution imaging of RNaseE and ribosomal proteins (Fig 5D) suggest that this energetic barrier exists for BR-bodies, as well (18).

Application of single-molecule tracking to less well characterized bacterial systems can confirm if LLPS is the assembly process or indicate alternative mechanisms. For example, McdB is known to undergo pH-dependent LLPS *in vitro* and interacts with carboxysome shell proteins (24, 95). Determining the diffusive behavior of McdB at or near carboxysomes will provide insight into its hypothesized LLPS activity *in vivo*. Furthermore, SMLM could support the model of LLPS-driven nucleoid compaction by Dps. Since RNAP maintains its ability to bind to its promoter under Dps-expressing conditions (47), single-molecule tracking could verify if RNAP has a slower, yet non-static, mobility in these condensed regions relative to cytoplasmic diffusion (16). Similarly, NusA and RpoC have different dynamics that correspond to free diffusion, binding (for NusA), and condensate confinement (23). To exclude the possibility of a transient DNA-binding mechanism (127), other transcription machinery and cytoplasmic proteins can be tracked.

Developing *in vivo* phase diagrams of phase-separating proteins will provide a clearer understanding of the phase transitions that occur within bacteria. Phase transitions are sensitive to environmental cues such as temperature, pH, and salt concentration, therefore knowing where phase boundaries exist in physical and chemical parameter space gives insight into the functional context of condensates (8). While microfluidic devices make the *in vitro* phase diagram accessible (132) *in vivo* characterization remains a challenge. It is possible to control protein concentrations to determine *c*_sat_, but detecting the formation of small condensates in bacteria requires super-resolution imaging and single-molecule sensitivity (8). Furthermore, component concentration in condensates can be estimated by using quantitative single-molecule protein counting techniques (133, 134). A uniform or random spatial distribution of components within a compartment can also provide a benchmark for LLPS (1). This quantification can be achieved using SMLM cluster analysis methods (135).

## Conclusions

The growing list of prokaryotic proteins that can phase separate indicates that LLPS may be a general mechanism by which bacteria spatially organize biochemical functions. While the surge in LLPS literature is exciting and creates a new field of study, strict experimental procedures based on a standard set of criteria (Fig 1) are necessary to definitively assign or discount LLPS. In this review, we described the criteria that have been developed based on years of characterization of condensates in eukaryotes. We also highlighted and discussed evidence for LLPS in bacteria to determine whether the data is conclusive. Furthermore, we commented on the general role of phase transitions in these organisms. Finally, we proposed various SMLM methods, which are ideally suited to overcome the challenges of size and dynamics, to examine and assess phase separation in small bacteria. Ultimately, the goal is to address each criterion for all hypothesized condensates to determine if LLPS is sufficient to explain the assembly of membraneless organelles and whether other criteria need to be evoked. Along these lines, a common theme across many of the proposed condensates is the observed selective boundary, which may provide a complementary criterion for assessing LLPS. We predict an expansion of the bacterial LLPS field and see a synergy between bacterial cell biology and super-resolution imaging as it pertains to phase separation.

## Author Contributions

All authors contributed to the writing and discussion of the manuscript.

## Acknowledgements

We thank Erin Goley and Saumya Saurabh for their critical reviews of the manuscript. J.S.B. was supported by the National Institute of General Medical Sciences (NIGMS) of the National Institutes of Health award number R21-GM128022. A.G.V. was supported by the National Science Foundation (CAREER Award No. 1941966). C.A.A. was supported by a University of Michigan Rackham Merit Fellowship.

